# Dietary and gut microbial variation among urban and rural populations of house mice (*Mus musculus domesticus*)

**DOI:** 10.64898/2026.04.30.721966

**Authors:** Samantha M. Giancarli, Adrienne E. Kasprowicz, Madeline Balman, René D. Clark, Stephen C. Kupchella, Logan J. Lacy, Andrew H. Moeller, Taichi A. Suzuki, Megan V. Phifer-Rixey

## Abstract

Urbanization can result in shifts in abiotic and biotic factors, including temperature, pollution, habitat type, pathogens, and diet, among others. These shifts can, in turn, shape the ecological and evolutionary trajectory of urban wildlife. The gut microbiota has the potential to mediate host-environment interactions, especially in the context of diet and disease, and thus may be a useful lens for understanding the impacts of urbanization. House mice (*Mus musculus domesticus*) are a cosmopolitan human commensal with a wealth of genomic and metagenomic resources. Here, we investigate patterns of variation in diet and gut microbial diversity, community composition, and function using a paired urban-rural sampling design in house mice from three metro regions in the eastern United States. First, using stable isotope analysis, we found that habitat—urban versus rural—was a major driver of variation in δ15N, suggesting a diet richer in animal proteins in cities. Next, using short-read sequencing of the 16S rRNA gene, we found that urban mice have lower gut microbial taxonomic diversity than their rural counterparts. We also found that community composition varied among urban and rural habitats, with differences largely reflecting shifts among closely related taxa. In particular, Prevotellaceae, a family known to be responsive to dietary quality, was differentially abundant, with lower abundance in urban habitats. Finally, we found differentiation in a few predicted microbial functions across habitat, primarily related to metabolism. Together, data across three independent sampling regions provide strong evidence that urbanization has the potential to shape the diet and the microbiome of house mice.

## 1. Introduction

Cities vary in many ways, including age, climate, and size. By definition, though, they are characterized by high-density human populations which are generally also accompanied by a common suite of habitat changes, including increased impervious surface area, heat, pollution, and waste (Bai et al., 2017; Seress & Liker, 2015). These changes have the potential to affect urban wildlife via processes such as habitat fragmentation, changes in resource availability, and the physiological and health consequences of light pollution, chemical and particulate pollutants (Bichet et al., 2013; Hofer et al., 2010; Kekkonen et al., 2011; M. Swaileh & Sansur, 2006), and thermal stress (Bateman et al., 2023; Blackwell, 2024; Frank & Backe, 2023). The gut microbiome can be an important mediator between a host and its environment, including processes such as metabolism, immunity, and homeostasis (Belkaid & Hand, 2014). For example, the gut microbiome can allow for plasticity and niche expansion, increasing a species’ fitness in a novel urban environment (Anders et al., 2022). Therefore, the gut microbiome may act as a valuable signal of how cities impact wildlife.

House mice (*Mus musculus*) offer a powerful opportunity to study the impact of urbanization in a globally distributed human commensal with extensive functional data, including many studies of the gut microbiome (Bär et al., 2020; Chevalier et al., 2015; Goertz et al., 2019; Moeller et al., 2019). Both environmental variables and dietary differences are expected to affect gut microbial variation among mice from urban and rural habitats. For example, the type of food available and access to food may differ among habitats. Cities may have more anthropogenic food and potentially more stochastic food access due to heavy human activity (Dillard et al., 2022; Sugden et al., 2020). On the other hand, rural populations of house mice are often associated with farms and other agricultural settings such that their diet may be dominated by animal feed. In addition, even if their diets are primarily human-derived in both habitats, human diets likely differ among rural and urban areas. In some cases, urban living is associated with reduced fiber content but more diverse foods and higher meat consumption (Cockx et al., 2018; Z. Yang et al., 2022). In other places, including the United States, cities have been linked to high-fat, high-protein diets (Gupta et al., 2017; Katsidzira et al., 2019). There is evidence that changes in human diets associated with urbanization can indirectly influence the diets of human commensals including rodents (Guiry & Buckley, 2018; Yu et al., 2017) along with their predators (Gámez et al., 2022; Hindmarch & Elliott, 2015). Diet shifts like those observed in cities may in turn affect the microbiome. Shifts in fiber (Morrison et al., 2020) and fat (Lang et al., 2018) in humans have been shown to alter diversity and composition of the gut microbiome (Azzam, 2021; Mancabelli et al., 2017; Paulo et al., 2023) and potentially make individuals more prone to disease (Zhang et al., 2025).

While there are now many studies that have investigated differences in the gut microbiome among urban and less developed areas in humans and other mammals (Anders et al., 2022; Bouilloud et al., 2024; Diaz et al., 2023; Giacomini et al., 2025; Heni et al., 2023; Lobato-Bailón et al., 2023; Łopucki et al., 2024; Schmidt et al., 2019; Stothart et al., 2019; Sugden et al., 2020; Wasimuddin et al., 2022), birds (Berlow et al., 2021; Dalziell & Welbergen, 2016; Gadau et al., 2019; Maraci et al., 2022; Murray et al., 2020; Schmitt et al., 2025; Solomon et al., 2023), and reptiles (French et al., 2018; Littleford-Colquhoun et al., 2019), most are limited to a single region, making it difficult to identify shared responses to urbanization. Moreover, no clear consensus has emerged (Nguyen et al., 2024). While some studies have found increased taxonomic diversity of the gut microbiota in urban animals (Berlow et al., 2021; Gurbanov et al., 2022; Phillips et al., 2018), others have seen reduced diversity in at least one metric (Maraci et al., 2022; Teyssier et al., 2020), and some saw no differences or no consistent pattern (Fuirst et al., 2018; Gadau et al., 2019; Stephens et al., 2021; Stothart et al., 2019; Teyssier et al., 2018). Again, most studies focus on a single region making it difficult to disentangle the effects of urbanization from other factors and/or include little functional data in which to contextualize the results (Nguyen et al., 2024). Systematic studies of the effect of urbanization on gut microbiome diversity and composition in house mice are limited. However, one study in an urban population of house mice suggests reduced diversity compared to other rodent species in less developed habitats (Bouilloud et al., 2024). Studies in rodents more broadly suggest shifts in diet and, in turn, altered gut microbiome community composition in cities (Anders et al., 2022).

Here, we characterized the gut microbiomes of wild house mice from three cities in the northeastern United States (Philadelphia, PA (PHL), New York City, NY (NYC), and Richmond, VA (RVA)), with each urban population matched with its surrounding rural counterpart. Importantly, this unique sampling design allowed us to test the generalizability of microbiome differences across habitat in three independent geographic locales. First, using stable isotope analysis, we found significant evidence that urban mice eat a higher trophic level than rural mice. This difference may reflect a diet with more human food and/or more insects or other live prey. Next, using 16S rRNA amplicon sequencing, we investigated how habitat (urban vs. rural) and other factors impact key aspects of gut microbiome composition and diversity. We found differences in taxonomic (alpha) diversity and community composition (beta diversity) between habitat types, namely that mice in urban areas had less diverse microbiomes. In addition, we identified specific taxa with ties to dietary shifts as differentially abundant among habitats. Finally, to better understand the functional consequences of those differences among urban and rural habitats, we applied the picrust2 (Douglas et al., 2020) pipeline, finding a handful of enriched and depleted pathways, mostly related to metabolism. Overall, our results suggest that urbanization impacts the microbiome in this human commensal, and likely as much or more so than geography.

## 2. Materials and Methods

### 2.1 Sampling

Using Sherman live traps baited with peanut butter and oats, we collected wild house mice from sites at least 500 m apart in three focal cities (New York City, NY (NYC), Philadelphia, PA (PHL), and Richmond, VA (RVA)) and in the rural areas surrounding them from June 2022 to May 2024 (Fig. 1; Supplementary Table 1; with approval from Drexel University IACUC Protocol No. LA-122-74 and Monmouth University IACUC Protocol No. Asp1802). Sample size varied (Supplementary Table 2), but at least eight individuals were collected for each habitat type within a metro region. We defined trapping areas as urban or rural based on the Global Human Settlement Layer (GHS-SMOD) settlement classification (Pesaresi et al., 2024; Schiavina et al., 2023). After trapping, mice were euthanized, sex was recorded, and ceca were dissected along with other tissues. Cecal contents were then either immediately stored at -80°C or briefly held on dry ice and then transferred to -80°C for storage.

**Figure 1.**
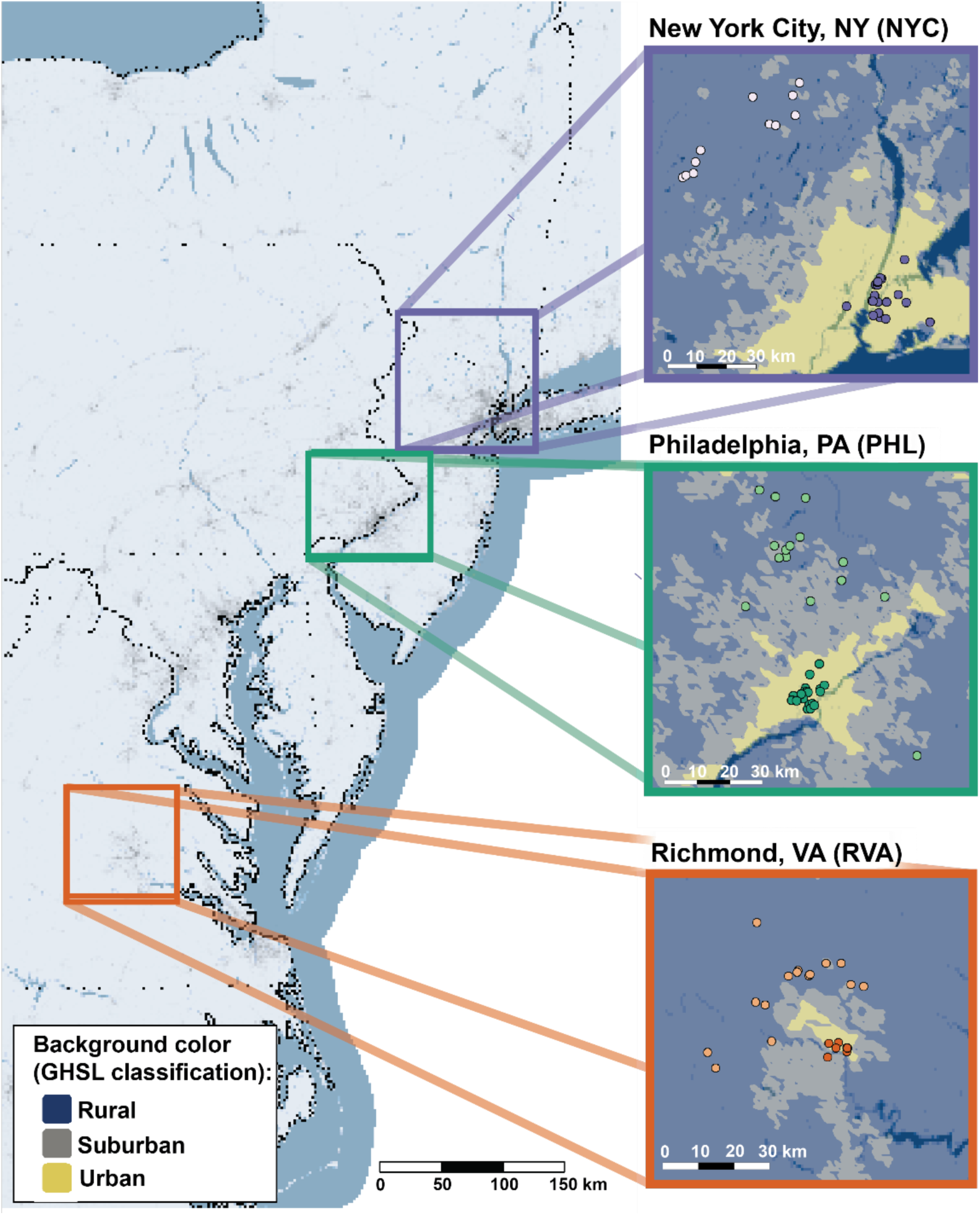
Mice were collected from highly developed and less developed sites in three metro regions: New York City, NY, Philadelphia, PA, and Richmond, VA. Inset background color indicates degree of urbanization (GHSL-SMOD (Pesaresi et al., 2024; Schiavina et al., 2023)). Each metro region inset is highlighted in a different color (NYC-purple; PHL-teal; RVA-orange). Lighter points within insets indicate locations designated as rural and darker points indicate urban sites.

### 2.2 Characterizing diet

Diet is a key determinant of microbiome variation. Because collection requires bait and mice have a short digestive passage time (Schwarz et al., 2002), we used stable isotope analysis of claws and fur for a coarse characterization of diet. From previous studies on other rodents, we expect the half-lives of stable isotopes in these tissues to reflect the host’s diet up to three months prior to capture (Kurle, 2009; Tieszen et al., 1983). The ratio of nitrogen-15 to nitrogen-14 (δ15N) can be used to make inferences regarding the trophic level of an animal’s diet. The heavier nitrogen isotope is more concentrated at higher trophic levels, thus higher δ15N suggests a diet richer in animal proteins (e.g., dairy or meat). Because C3 and C4 plants fix carbon isotopes at different rates, the ratio of carbon-13 to carbon-12 (δ13C) can be used to assess the relative contribution of C4 plants like corn vs C3 plants like wheat and oat to diet (Hobson, 2023). Therefore, higher values of δ13C reflect a diet richer in C4 plants. Claw clippings were collected from the left forelimb. In some cases, clippings from the left forelimb were insufficient for analysis and were augmented by hindlimb clippings (n=19). Because different tissues may reflect slightly different timescales and data estimating such timescales for house mice are not available, we also tested fur. We collected fur from the right ventral side of the prepared museum specimens (n=90; Supplementary Table 2). Claw clippings and fur were then rinsed with DI water and left to dry. A Thermo Scientific Delta V continuous flow mass spectrometer was used to assess δ15N and δ13C (Center for Stable Isotopes at the University of New Mexico). To investigate drivers of variation in these ratios, we performed Analyses of Variance (ANOVA) regressing δ15N and δ13C respectively against habitat, metro region, and their interaction. All statistical analyses were performed using RStudio (version 2023.06.0+421).

### 2.3 DNA extraction, sequencing, and bioinformatics

We extracted DNA from the cecal contents of each wild mouse using the QIAGEN PowerFecalPro kit. Amplicon libraries of the V3 and V4 regions of the 16S rRNA gene were prepared using the Zymo Quick-16S Plus NGS Library Prep kit and then sequenced on the Illumina NextSeq2000 platform using a 600-cycle flow cell kit (SeqCoast Genomics, Portsmouth, NH). Reads (300 bp paired-end) were demultiplexed and trimmed using DRAGEN v4.2.7. We then analyzed the resulting FASTQ files using the amplicon version of QIIME2 (version 2024.5). Specifically, reads were filtered requiring quality scores ≥ 20 and the Deblur function was used to denoise the sequences and create a table of representative amplicon sequence variants (ASVs; Supplementary File X). After these steps, there were 90 samples from unique locations with data of sufficient quality for analysis (Supplementary Table 2). We used the pretrained SILVA 138.1 naïve Bayes human stool weighted feature classifier formatted for use in QIIME2 to categorize each ASV into taxonomic groups (Ii et al., 2021; Robeson et al., 2020). The feature classifier reliably classified the ASVs into domains, phyla, classes, and orders. However, there were several ASVs for which genus was not resolved, and, in a few cases, SILVA was unable to resolve classification at the family level. As a result, we opted to perform analyses of microbial community at the ASV level and to perform differential abundance analyses at multiple levels (family, genus, and ASV). We then rarefied a dataset to a depth of 5000 reads in QIIME2, which was used for all analyses except differential abundance for which raw count data were used.

### 2.4 Diversity Analyses

Using the phyloseq R package (McMurdie & Holmes, 2013), we joined sample metadata with the QIIME2-generated datasets (including a phylogeny of ASVs, abundance tables, and taxon-assignment tables) for phylogenetically informed microbial community analyses. Using phyloseq’s *estimate_richness* function, we then compared taxonomic richness (alpha diversity) across habitats (urban, rural) and metro regions (NYC, PHL, RVA) using four measures: observed ASV richness, Shannon diversity, Simpson diversity, and Faith’s Phylogenetic Diversity (PD). The Shannon Index reflects evenness while the Simpson Index is driven by the abundance of common taxa. Faith’s PD is a measure of diversity that integrates phylogenetic distance between observed features. The contributions of metadata (e.g., habitat, metro region, sex, stable isotope ratios, and their interactions) to variation in alpha diversity were tested via ANOVA, Tukey’s HSD, and linear mixed-effects models with the general formula: [alpha_diversity metric] ∼ habitat + sex + 1|metro_region.

Beta diversity refers to variation in the gut microbial community assemblage among samples. Measures of beta diversity can integrate data regarding the presence/absence and/or abundance of taxa/ASVs. We first constructed a matrix of Bray–Curtis pairwise distance (BCD) among individuals (Bray & Curtis, 1957). For comparison, we also constructed matrices of Jaccard distance (Jaccard, 1912) and weighted and unweighted UniFrac distance (Lozupone et al., 2011). BCD integrates both presence/absence and abundance data. Jaccard distance is constructed solely based on the presence/absence data. Unweighted UniFrac distance uses presence/absence data as well as the phylogenetic relatedness among the bacterial taxa. Weighted UniFrac extends this by weighting the data based on abundance. We then performed a principal coordinate analysis (PCoA) of BCD to visualize overall patterns in beta diversity with both the full dataset and then individually for each city. Next, associations between host gut microbial assemblage (distance matrices as described above) and factors (e.g., habitat, metro region, sex, and their interactions) were tested by performing a permutational analysis of variance (PERMANOVA) using the R function *adonis2* within the vegan package (Jari et al., 2001) and using the general formula: [distance matrix] ∼ habitat x region x sex x δ13C x δ15N. Finally, the pairwise_adonis function was then used to test for differences among levels of individual factors (Arbizu, 2020).

### 2.5 Differential Abundance Analyses

We used Analysis of Compositions of Microbiomes with Bias Correction 2 (ANCOM-BC2) (Lin & Peddada, 2020), to identify differentially abundant taxa. ANCOM-BC2 implements sample-specific terms to address differences in sampling depth and fits log-linear models to calculate differential abundance of bacterial taxa between groups of samples. Additionally, it compares the counts of each taxon in all samples and iteratively fits a model to infer a factor accounting for compositional bias (i.e., the sum of relative abundances of taxa in each sample being equal to 1). This factor then gets incorporated into the prediction of the effects of categorical metadata in differential abundance analyses (Lin & Peddada, 2020). Compared to other approaches, ANCOM-BC2 has a lower false discovery rate (Pelto et al., 2025). We constructed models at the level of family, genus, and ASV respectively. Using the ANCOM-BC2 R package, we regressed the assemblage of gut microbes against both habitat, metro region, and their interaction with “rural” and “NYC” being the reference groups against which the other habitat and metro regions are compared. P-values were adjusted to reduce the false discovery rate (Benjamini & Hochberg, 1995), and a threshold of 0.1 for significance was used. To avoid structural zeros, we implemented a pseudo-count of 0.001, as well as a pseudo-sensitivity test built into the ancombc2 function to avoid false discovery resulting from adding a pseudo-count.

### 2.6 Inferring functions of gut microbial taxa

To explore the functions of gut microbial taxa in the samples, we implemented the picrust2 pipeline (Douglas et al., 2020) using our unrarefied count data and representative sequences for each ASV. This pipeline is designed to predict functions associated with taxa using the following databases of metabolic pathways: MetaCyc, the Kyoto Encyclopedia of Genes and Genomes (KEGG), and the Enzyme Commission (EC). We then used the ggpicrust2 package in R (C. Yang et al., 2023). to identify differentially abundant functions between habitats. The ggpicrust2 package uses both abundance data and predicted functions from each ASV, groups similar functions, and then implements differential representation analysis and principal components analysis (PCA). We used the method “LinDA” within ggpicrust2 as this has been shown to reliably assess differential abundance in compositional data (C. Yang et al., 2023). We also used PERMANOVA analyses to compare overall pathway representation between metadata categories, regressing distance matrices constructed from feature tables of metabolic pathways using the general formula and using the general formula: [distance matrix] ∼ habitat x region x sex. Finally, we considered functional diversity and redundancy in samples, characterized here as the count of inferred KEGG orthologs and KEGG ortholog count divided by Faith’s PD, respectively. These two terms were each regressed in ANOVAs against habitat, metro region, sex, and their interactions.

## 3. Results

### 3.1 Urban mice eat at higher trophic levels than rural mice

Habitat, but no other terms, contributed significantly to variation in δ15N in both fur (p = 8.92 x 10^-4^, F = 11.94, η^2^ = 0.13) and claws (p = 2.41 x 10^-5^, F = 20.22, 17^2^ = 0.21; Figs. 2, S1, S2; Supplementary Table 3). Using Tukey’s HSD, we found that δ15N was higher in urban mice compared to rural mice in fur (Figs. 2 & S2; Supplementary Table 4) and claws (Figs. S1 & S2; Supplementary Table 3) suggesting that urban mice eat at higher trophic levels. This difference could be driven by more protein from human food sources and/or more insects or other live prey in their diet. Tukey’s HSD indicated significant effects of the habitat-sex interaction in δ15N in both tissues, as well as effects of habitat-region and habitat-region-sex interactions for δ15N in claws (p-adjusted <0.05; Supplementary Table 4). There was no evidence that any of the included factors contributed to variation in δ13C in claws or fur (Supplementary Tables 3 and 4).

**Figure 2.**
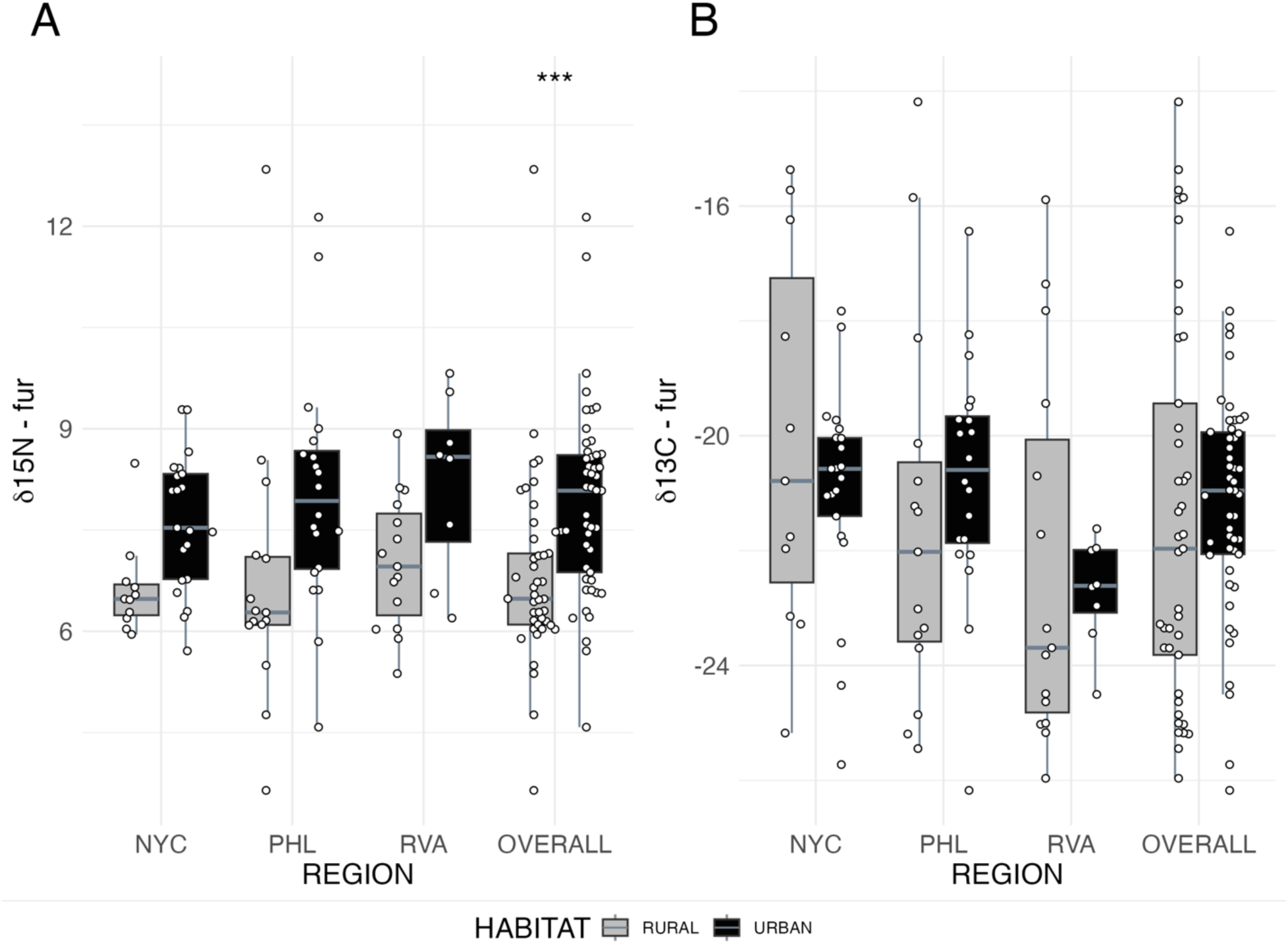
Habitat contributed to variation in A) δ15N but not B) δ13C in fur (for claws, see Fig S1; NYC=New York City, PHL = Philadelphia, RVA=Richmond). Each point represents one gut sample from one mouse. Box colors represent habitat type (rural = gray, urban = black). *** p < 0.001.

### 3.2 Taxonomic diversity is higher in rural mice compared to urban mice

Results of ANOVA indicated that habitat contributed significantly to variation in ASV richness (p=4.31×10^-2^, F = 4.34, η^2^ = 0.09) and Faith’s PD (p=2.85×10^-2^, F = 5.13, 17^2^ = 0.10; Fig. 3, S3, Supplementary Tables 5). Tukey’s HSD showed that rural samples have significantly higher ASV richness and PD than urban samples (Supplementary Table 6). While these tests did not identify a significant contribution of habitat to variation in Shannon and Simpson Diversity, trends were consistent with results from ASV richness and Faith’s PD, with higher diversity in rural populations (Fig. 3, Supplementary Tables 5 & 6). Moreover, there was a significant contribution from the interaction between habitat and sex to both Shannon (p = 2.09 x 10^-2^, F = 5.74, 17^2^ = 0.12) and Simpson indices (p = 3.22 x 10^-2^, F = 4.89, 17^2^ = 0.10; Supplementary Table 5) suggesting that the effects of habitat are contingent on sex. Male house mice are expected to disperse more than female mice (Pocock et al. 2005) which may contribute to this pattern (Fig. S3). In none of these ANOVAs were any other top-level factors significant. However, we did detect significant contributions from the interactions between sex and both stable isotope ratios to Simpson diversity (p = 4.89 x 10^-2^, F = 4.10, 17^2^ = 0.09; Supplementary Table 5). The linear mixed-effect model using metro region as a random effect did not detect a significant effect of region on the effects of habitat and sex for any metric of alpha diversity (Supplementary Table 7).

**Figure 3:**
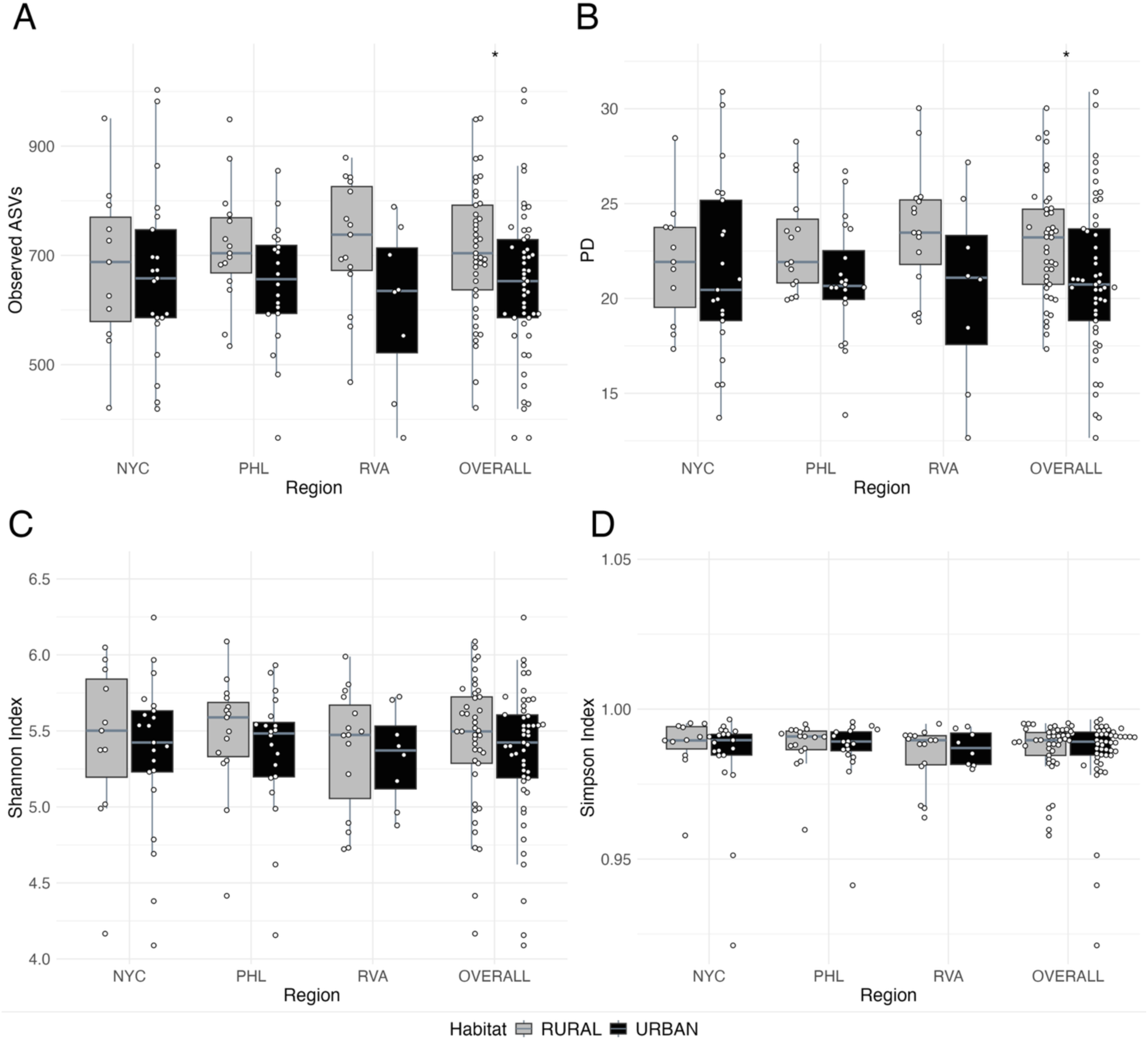
Rural populations are significantly more diverse as measured by A) observed ASVs (p=4.31×10^-2^; R^2^=0.09) and B) Faith’s PD (p=2.85×10^-2^; R^2^=0.10) but not using C) Shannon Index (p=4.65×10^-1^; R^2^=0.01) and D) Simpson Index (p=7.97×10^-1^; R^2^=0.01). Each point represents one gut sample from one mouse. Box colors represent habitat type (rural = gray; urban = black). * p < 0.05.

### 3.3 Habitat is a primary driver of variation in gut microbial community composition

A PERMANOVA model regressing BCD on habitat, metro region, sex, stable isotope ratios from fur, and their interactions found significant effects of habitat (p = 2.0×10^-3^; F = 1.77; R^2^ = 0.02), metro region (p = 3.0×10^-3^; F = 1.32; R^2^ = 0.03), and the interaction between δ13C and δ15N (p= 4.2×10^-2^ ; F = 1.23; R^2^=0.01; Table 1; Supplementary Table 8). Repeating the PERMANOVA using stable isotope ratios from claw clippings yielded similar results with significant effects of habitat (p=1.0×10^-3^; F = 1.77; R^2^=0.02) and metro region (p=3.0×10^-3^; F = 1.28; R^2^=0.03; Supplementary Table 8). Pairwise PERMANOVA indicated a significant difference between the gut microbial communities of rural and urban mice (p=3.4×10^-2^; F = 2.23; R^2^=0.02; Supplementary Table 9), but not among samples of different metro regions or sexes. PCoA on BCD showed little separation between habitats along the first and second axes (Fig. 4A). However, data separated across habitats on the third axis (Fig. 4B).

**Figure 4.**
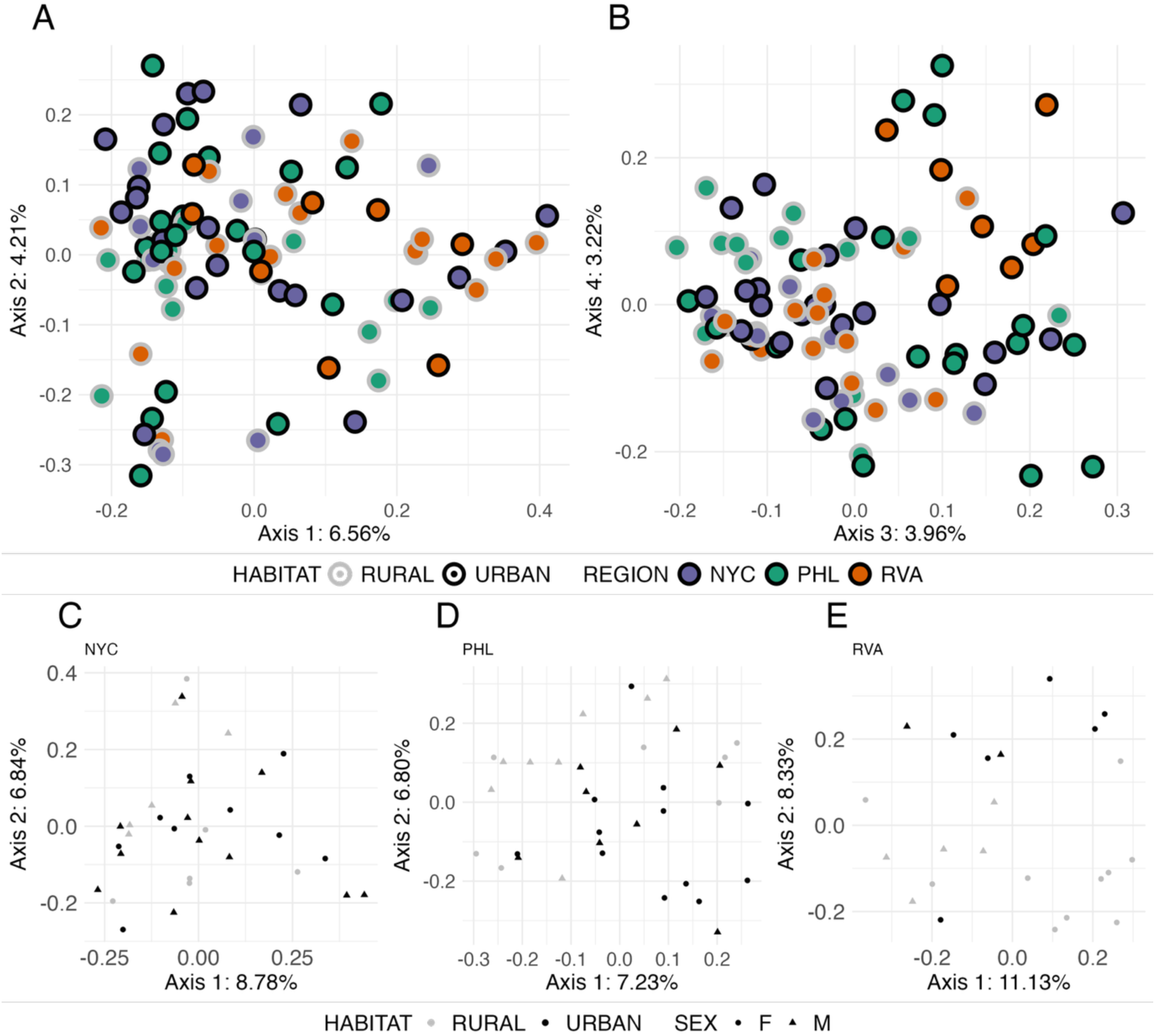
Habitat contributes to variation in community composition (p=0.02; R^2^=0.02). PCoA on Bray–Curtis Distance on A) Axes 1 and 2 and B) Axes 3 and 4 which shows separation between habitats. Habitat is indicated by circle outline color (rural = gray, urban = black) and metro region is indicated by circle fill color (NYC = purple, PHL = green, RVA = orange). C-E) Axes 1 and 2 of PCoAs of Bray–Curtis Distance specific to the C) New York, D) Philadelphia, and E) Richmond metro regions. Each point represents one mouse gut sample. Point colors represent habitat type (rural = gray; urban = black), and shape represents host sex (circle = female; triangle = male).

The same analyses were then performed on Jaccard, Unweighted UniFrac, and Weighted UniFrac distance (Figs. S4, S5). We found evidence that habitat (p = 1.0×10^-3^; F = 1.41; R^2^=0.02), metro region (p = 6.0×10^-3^; F = 1.17; R^2^=0.03), the interaction between habitat and metro region (p = 4.5×10^-2^; F = 1.10; R^2^=0.02), and the interaction between δ13C and δ15N (p = 4.6×10^-2^; F = 1.13; R^2^=0.01) contributed to variation in Jaccard distance (Table 1, Supplementary Table 8). Pairwise PERMANOVA tests indicated habitat-based differences (p = 2.0×10^-3^; F = 1.41; R^2^=0.02; Supplementary Table 9) and metro region-based differences (PHL vs. RVA: p = 1.5×10^-2^; F = 1.26; R^2^=0.02, and RVA vs. NYC: p = 4.8×10^-2^, F = 1.23; R^2^=0.02). Similar to BCD, differences between habitats and metro regions could be seen on the third axis of the PCoA (Fig. S5).

Using Unweighted UniFrac distance, we found similar results to those with BCD and Jaccard distances. A PERMANOVA identified habitat– (p=1.0×10^-3^; F = 2.14; R^2^=0.02) and metro region–driven (p = 8.0×10^-3^; F = 1.21; R^2^ = 0.03) variation in beta diversity, as well as variation driven by their interaction (p = 1.2×10^-2^; F = 1.17; R^2^ = 0.03). In addition, the interactions between habitat and sex (p = 6.0×10^-3^; F = 1.29 R^2^ = 0.01;) and metro region and sex (p=3.0×10^-3^; F = 1.20; R^2^=0.03; Table 1; Supplementary Table 8) contributed significantly to variation. Pairwise PERMANOVA identified significant habitat-based differences (p = 1.0×10^-3^; F = 2.09; R^2^=0.02) and between PHL and RVA (p = 4.5×10^-2^; F = 1.23; R^2^ = 0.02; Supplementary Table 9). As with Bray–Curtis and Jaccard distances, a PCoA on Unweighted UniFrac distance between samples showed a separation between rural and urban samples along the third axis (Fig. S5). Like its unweighted counterpart, Weighted UniFrac distance is based on the relatedness among taxa, but it also weights distances based on the abundances of those taxa. For this measure of between-sample distance, there was no evidence that any of the factors or their interactions contributed to variation in beta diversity (Table 1, Supplementary Tables 8 & 9, Fig. S5).

Because metro region contributed significantly to variation in beta diversity, we also considered the effects of habitat, sex, and stable isotope ratios on BCD within metro regions. Notably, PCoAs consistently showed clear separation of urban and rural samples (Figs. 4, S8) along either the first or second axis in all three metro regions. Habitat contributed significantly to variation in BCD in all three PERMANOVA analyses (NYC: p = 1.5×10^-2^; F = 1.34; R^2^=0.04; PHL: p = 3.0×10^-3^; F = 1.50; R^2^ = 0.04; RVA: p = 1.9×10^-2^; F = 1.35; R^2^=0.06; Supplementary Table 10).

PERMANOVAs also identified significant contributions to variation in the NYC metro region from the interactions between sex and δ13C, habitat and δ15N, and the between sex and both stable isotope ratios (Supplementary Table 10).

We also asked whether metro region, sex, and stable isotope ratios and their interactions contributed to variation in BCD among only rural and only urban populations respectively. No factors were identified as significant contributors to variation in the PERMANOVA of only rural populations (Supplementary Tables 12 & 13). However, the urban-specific PERMANOVA identified metro region (p = 3.0×10^3^; F = 1.34; R^2^ = 0.05) and the interaction between metro region and δ15N (p = 3.2×10^2^; F = 1.19; R^2^ = 0.05; Supplementary Table 12). Pairwise PERMANOVA indicated a significant difference between the bacterial communities of urban PHL and urban RVA samples (p-adjusted = 4.8 ×10^2^; F = 2.37; R^2^ = 0.08; Supplementary Table 13).

### 3.3 Taxa functionally linked to diet are differentially abundant across habitat

ANCOM-BC2 identified one family as differentially abundant, Prevotellaceae, which is underrepresented in urban habitats (p-adjusted=1.90 x 10^-5^, Supplementary Table 14). Bacteria within Prevotellaceae are anaerobes that are prevalent in the gut microbiota of humans, sheep, and cattle, where they help to break down proteins, carbohydrates, and fiber. Genus-level analyses identified three differentially abundant genera. UCG-001, a genus within Prevotellaceae, which was underrepresented in urban habitats (p-adjusted = 4.4 x 10^-7^), *Negativibacillus* which was overrepresented in urban habitats (p-adjusted = 3.85 x 10^-8^, Supplementary Table 14). *Negativibacillus* falls within the family Ruminococcaceae and has been putatively associated with lower quality (Balasubramaniam and Srinivasan 2024) and higher protein diets (Singh et al. 2025; Segev et al. 2026). The genus *Anaerovorax* is also overrepresented in urban hosts, and is associated with the fermentation of polyamines into short-chain fatty acids such as butyrate (Matthies et al., 2000). Butyrate is known to be beneficial in the face of insulin resistance and high-fat diet-induced obesity (Gao et al., 2009; Henagan et al., 2015). At the ASV level, our analysis identified one differentially abundant taxon in the dataset, a member of Lachnospiraceae, that is overrepresented in urban habitats (p-adjusted = 3.43 x 10^-3^, Supplementary Table 14). Members of Lachnospiraceae are also known producers of short-chain fatty acids (Vacca et al., 2020a). To test for parallel instances of differential abundance of taxa in each metro region, we performed ANCOM-BC2 analyses at the same three taxonomic levels on the samples in each metro region individually. Consistent with the overall analysis, we found that Prevotellaceae is underrepresented in urban NYC mice, and the genus UCG-001 within Prevotellaceae is underrepresented in urban mice in both NYC and PHL.

Differential abundance analysis using ggpicrust2 and the MetaCyc metabolic pathway data suggested that three predicted pathways were underrepresented in urban habitats (p-adjusted <0.05; Supplementary Table 15, Fig. S9; see Supplementary Analyses). These predicted pathways were associated with coenzyme factor 420 biosynthesis (involved in redox reactions in several bacterial taxa, including those for antibiotic production and xenobiotic biodegradation; Grinter & Greening, 2021), the biosynthesis of CDP-glucose-derived O-antigen building blocks (linked to antibiotic resistance and host–microbe interactions in gram-negative bacteria (Lerouge & Vanderleyden, 2002)) and the degradation of mandelate into acetyl-CoA. Mandelate and its derivatives are widely used in the pharmaceutical industry and have antimicrobial properties (Choińska et al., 2021).

We identified 49 significantly differentially represented KEGG Orthologs (KOs) between rural and urban habitats (p-adjusted <0.05; Supplementary Table 15, Fig. S9). These KOs are mainly associated with antimicrobial resistance, bacterial signaling and quorum sensing, metabolism and biosynthesis of amino acids, sugars and nucleic acids, and the cell cycle. CDP-4-dehydro-6-deoxyglucose reductase, E3 (K00523) was underrepresented in urban habitats. Similar to a pathway identified using MetaCyc, this ortholog is involved in the biosynthesis of components of O-antigens (Liu & Thorson, 1994). Only one KO, arginine decarboxylase (K02626), was overrepresented in urban habitats. This ortholog is key in the metabolism of arginine and proline, as well as the biosynthesis of polyamines, a multipurpose class of molecules that is shown to have beneficial effects on metabolic processes in mouse models (Ramos-Molina et al., 2019).

Only one pathway from the Enzyme Commission database was identified by LinDA as significantly differentially represented across habitats: CDP-4-dehydro-6-deoxyglucose reductase was found to be underrepresented in urban habitats (p-adjusted=9.61×10^-3^; Supplementary Table 15, Fig. S9). This enzyme works within the aforementioned super-pathway of CDP-glucose-derived O-antigen building blocks biosynthesis (Schomburg & Stephan, 1994).

## 4 Discussion

### 4.1 Growing evidence for dietary shifts to higher trophic levels in cities

Our results point to consistent shifts in diet across habitat with mice in urban areas eating at a higher trophic level. This difference suggests a greater reliance on anthropogenic food sources, companion animal food sources, and/or live prey in urban mice. The presumed diets of rural mice, which were mostly caught in agricultural settings, in contrast, were likely made up of primarily horse and livestock feed. Many such feeds are plant-based, such that opportunistic diets of anthropogenic food and pet food in cities would likely contain more animal-based protein. Consistent with our findings, lower δ15N has previously been found in house mice with access to livestock feed, as opposed to anthropogenic sources (Balčiauskas et al., 2023). An increase in δ15N has also been reported in urban populations of other rodents (Anders et al., 2022; Guiry & Buckley, 2018; Ouellette & Schulte-Hostedde, 2025) and lizards (Littleford-Colquhoun et al., 2019) compared to their respective rural counterparts. Anders et al. attribute the urban-associated increase in δ15N to expansions of the dietary niches of both rodent species in their study (Anders et al., 2022).

The carnivore-to-herbivore dietary axis has been shown to shape gut microbiota in mammals (Cheng et al., 2025; Ley et al., 2008; Muegge et al., 2011; Zhu et al., 2018; Zoelzer et al., 2021). For example, both within humans and mice, abundance and/or dominance of taxa within *Bacteroides* and *Prevotella* are seen to covary with diet (Gorvitovskaia et al., 2016, 2016; Kovatcheva-Datchary et al., 2019; Wu et al., 2011). *Bacteroides* are overrepresented in hosts eating more animal fat and protein and *Prevotella* are enriched in hosts eating more plant-based diets (De Filippo et al., 2010; Wu et al., 2011; Yatsunenko et al., 2012). While we did not observe that stable isotope ratios contributed significantly to overall variation in measures of alpha and beta diversity, differential abundance analyses did identify shifts in Prevotellaceae across habitat. Prevotellaceae was underrepresented in urban habitats, where we also found evidence that mice were eating at higher trophic levels. In addition, *Negativibacillus* was overrepresented in urban habitats. In a recent human study, *Negativibacillus* was a predictor of meat consumption and was positively correlated with fast food consumption and negatively correlated with indicators of a high-quality diet (Singh et al., 2025). Data from The Human Phenotype Project also show that *Negativibacillus* is enriched in carnivore diets and depleted in vegetarian and pescatarian diets (Segev et al., 2026). Therefore, the observed differences in Prevotellaceae and *Negativibacillus* abundance are consistent with the dietary differentiation we observed and mirror patterns in humans, suggesting a shift towards a more anthropogenic diet in cities.

### 4.2 Evidence for the impacts of urbanization on gut microbiome diversity and composition

We found consistent evidence that alpha diversity of the gut microbiome is lower in urban mice. Moreover, no other factors were identified as significant drivers of taxonomic diversity. In addition, we found that habitat contributed to variation in beta diversity both overall and within individual metro regions. Evidence across multiple metro regions makes it less likely that this difference is idiosyncratic or due to other factors. Importantly, in our models, effect sizes for habitat were similar to effect sizes for metro region, which is expected to be a major driver of microbiome variation (Goertz et al., 2019; Linnenbrink et al., 2013; Mobeen et al., 2018). There is no clear consensus yet on how urbanization impacts gut microbiome diversity and composition, even within rodents. Studies like this one that consider patterns across cities are needed in more taxa with diverse life histories to help identify emergent patterns and the factors that drive them.

It is important to note that in contrast to other measures of beta diversity, habitat did not contribute significantly to variation in Weighted UniFrac distance. This measure accounts for both abundance and phylogenetic distance among ASVs. Therefore, these results suggest that the rural-urban compositional differences captured by other methods may be driven by bacteria that are closely related and rare among the communities. For example, rare ASVs in the gut of rural mice that are absent in urban mice could have taxonomically similar counterparts in urban mice. This hypothesis would predict relatively smaller differences in functionality of bacterial communities, which is consistent with our results. A study of three murid species classified as “urban adapters” (Bouilloud et al., 2024) also found few differences in gut microbial community composition along a rural-urban gradient, putatively attributing those results to functional redundancy in those communities along the gradient (Bouilloud et al., 2024). In fact, some argue that functional redundancy in the gut microbiota may facilitate mediation of response to environmental disturbance (Moya & Ferrer, 2016).

### 4.3 The functional consequences of urbanization on the gut microbiome

Overall, we found evidence of differentially abundant taxa and of differential representation of predicted microbial functions and pathways between habitats that are of potential functional significance in urban contexts. Notably, the family Prevotellaceae and the genus UCG-001 contained within it are underrepresented in the guts of urban mice and the genus *Negativibacillus* is overrepresented. As discussed above, these taxa have been linked to dietary differences in protein and overall quality in humans (Segev et al., 2026; Singh et al., 2025; Wu et al., 2011). These results underscore the importance of diet in microbiome composition and function and also provide evidence of a shared response to dietary shifts in humans and wild house mouse populations. While consensus on the impacts of urbanization on microbiome diversity and composition remains limited, results like these suggest testable hypotheses for studies in other mammalian taxa.

In addition, we also found overrepresentation of the genus *Anaerovorax* and an ASV in Lachnospiraceae in urban hosts. Both of these taxa are related to metabolism and diet and in particular production of short-chain fatty acids (SFCA) (Matthies et al., 2000; Vacca et al., 2020b). SFCA are diverse in function, serving as energy sources for intestinal tissues, regulators of innate immune cells, raw materials for sugar and lipid catabolism, and more (Corrêa-Oliveira et al., 2016; Yao et al., 2022). While our study does not address the specific mechanisms driving shifts in these taxa, results suggest that diet has large role in mediating the impacts of urbanization on the gut microbiome. Moreover, differences in diet that we could not assess with stable isotopes alone may be contributing to the habitat-driven effects on the microbiome that we uncovered.

There was also evidence of underrepresentation of predicted enzymes and pathways associated with building O-antigens in the guts of urban mice. An underrepresentation of O-antigen building pathways has also been reported in mourning doves fed anthropogenic food (Mohr et al., 2022). O-antigens are important in mediating host immune response in gram-negative bacteria (Lerouge & Vanderleyden, 2002), suggesting some differences in host-microbial interactions among habitats. In addition, a number of other pathways are underrepresented in urban mice that are related to bacterial signaling and quorum sensing, metabolism and biosynthesis of amino acids, sugars and nucleic acids, and the cell cycle. Nevertheless, we did not find that habitat contributed to variation in overall functional diversity or functional redundancy. It is worth noting, however, that inferring gut microbial metabolic functions using only regions of the 16S rRNA gene, as we did here, can be less reliable than analyses using higher resolution data generated by metagenomics, metabolomics, and meta-transcriptomics (Matchado et al., 2024).

## Conclusion

Here, we compared the diets and gut microbiota of urban and rural house mice in three metro regions. House mice are human commensals, and thus there was no clear *a priori* prediction of how much urbanization would impact diet or the gut microbiome. Nevertheless, we found significant evidence of differences in dietary trophic level and consistent evidence of significant impacts of habitat on diversity and composition of the gut microbiota. These effects were of similar or greater impact than the effects of metro region and diet as measured with stable isotopes. Differentially represented taxa and pathways across habitats related to metabolism and immunity and, in some cases, mirror patterns in humans. Together, these results highlight the potential for urbanization to shape diets and microbiomes, even in a human commensal, and, in turn, impact functions key to host fitness. This study provides a firm foundation for ongoing comparative work in other cities and taxa.

## Data Availability

Raw sequencing data are available via the Short Read Archive (NCBI BioProject# PRJNA1460303). All code is available via Github (https://github.com/PhiferrixeyLab/giancarli_2026).

## Supporting information

Supplemental_Figures_and_Analyses

## Acknowledgements

This research was supported by NSF Division of Environmental Biology CAREER Award #2332998 to Dr. Megan Phifer-Rixey. Dr. Taichi A. Suzuki is supported by the National Institutes of Health (R35 GM160076). Dr. Andrew H. Moeller is supported by the National Institutes of Health (R35 GM138284 and R01 DK139214). We would like to thank Erin Oscar, Teresa Pacelli, and Jason Munshi-South for their assistance with sampling. We also thank Nina Mallalieu, Zade Alafranji, Hannah Graham, and Jessica Bonnen for their helpful feedback and the students of Drexel Biology’s T380 research in evolutionary genomics course for their input. This work used the San Diego Super Computer Center through allocations BIO240194 and BIO230113 from the Advanced Cyberinfrastructure Coordination Ecosystem: Services & Support (ACCESS) program, which is supported by U.S. National Science Foundation grants #2138259, #2138286, #2138307, #2137603, and #2138296. We wholeheartedly thank all of the participants who welcomed our project into their communities in Connecticut, New Jersey, New York, Pennsylvania, and Virginia and made this research possible.

