## Supplemental_Figures_and_Analyses for "Dietary and gut microbial variation among urban and rural populations of house mice (*Mus musculus domesticus*)"

**Supplementary Figures**

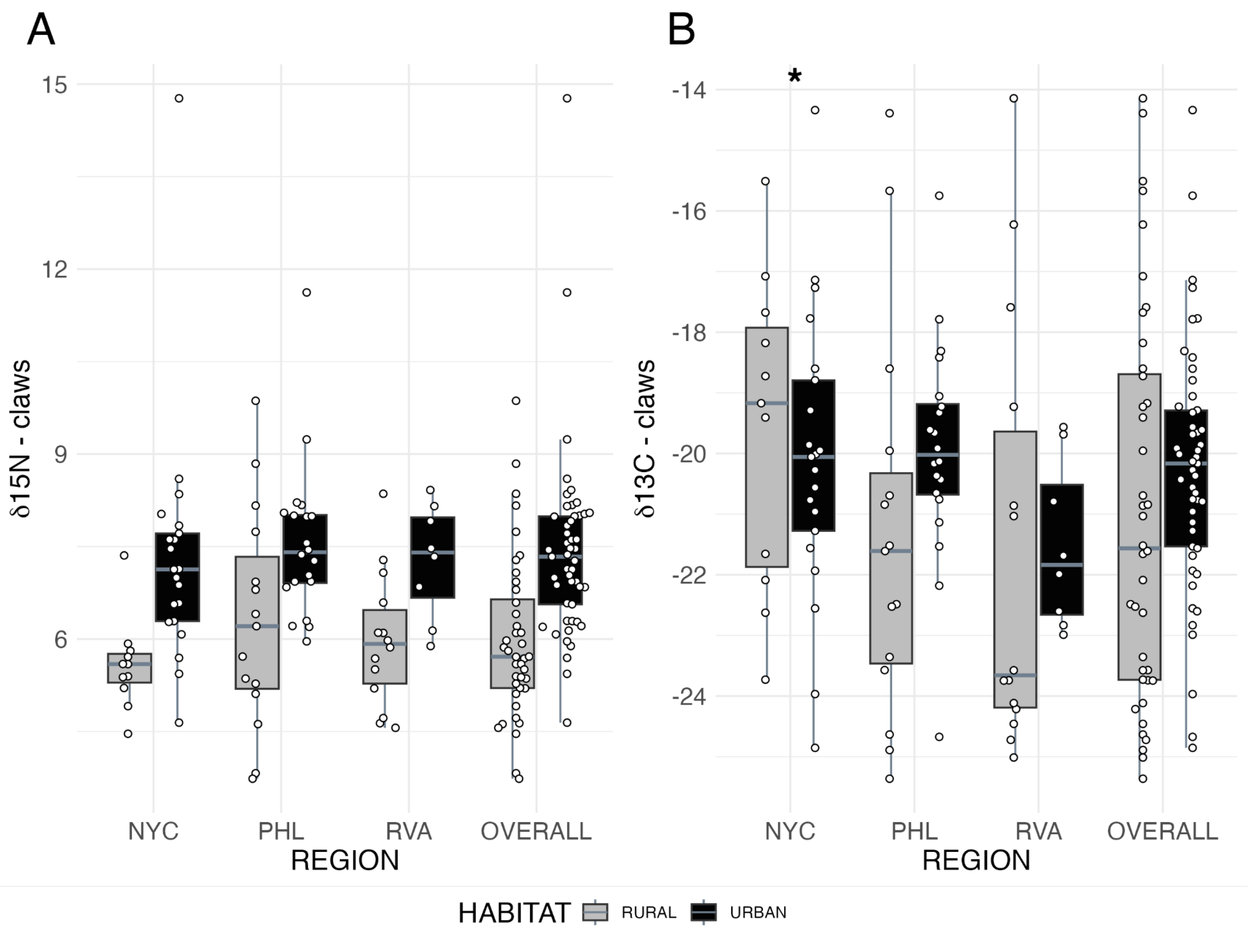


Figure S1. Boxplots of A. nitrogen and B. carbon stable isotope ratios from claws of host mice separated by metro region. Colors correspond to habitat types.

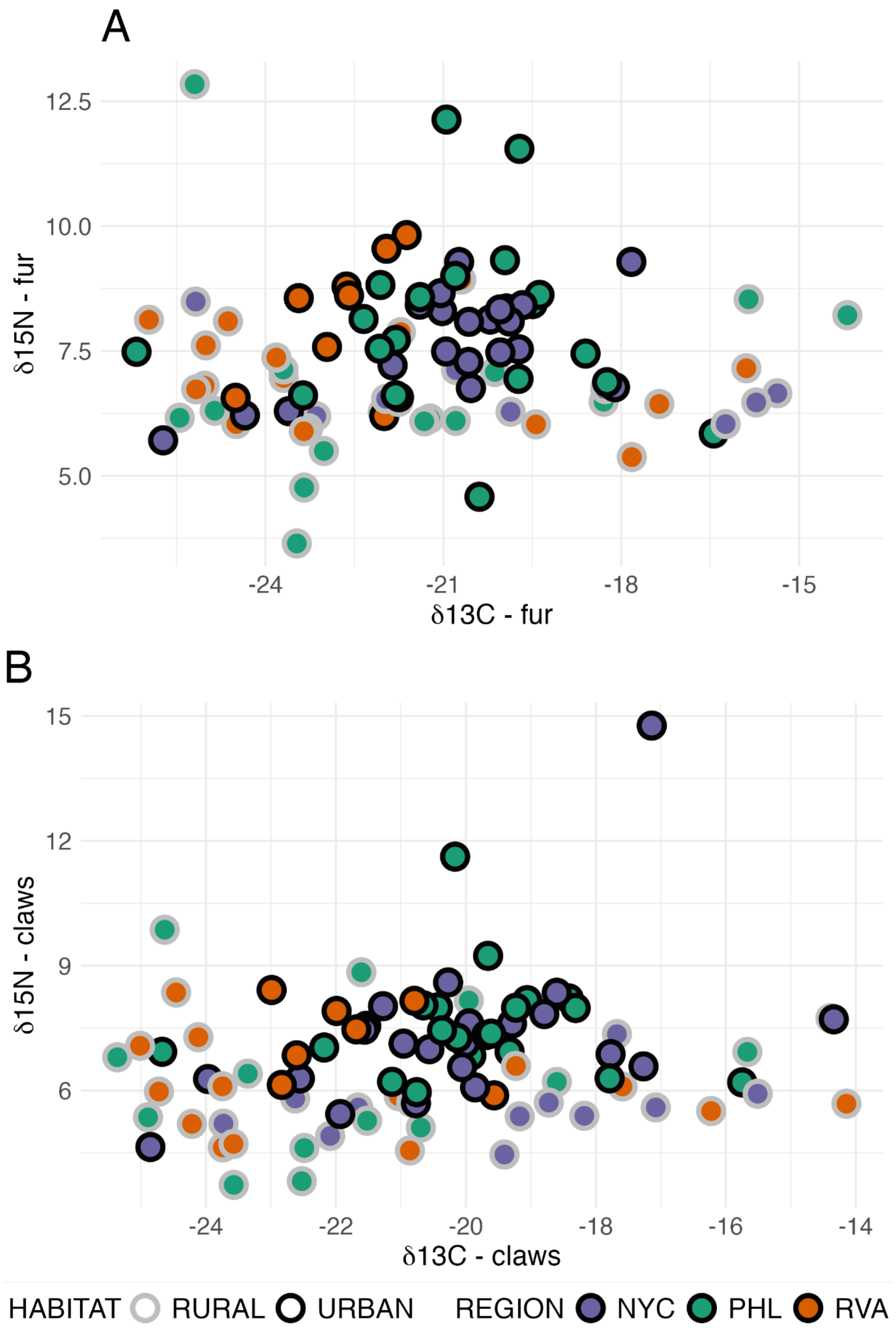


Figure S2. Nitrogen vs. carbon stable isotope ratios from A. fur and B. claws of host mice. The outer color of each circle indicates habitat type and the inner color corresponds with the metro region.

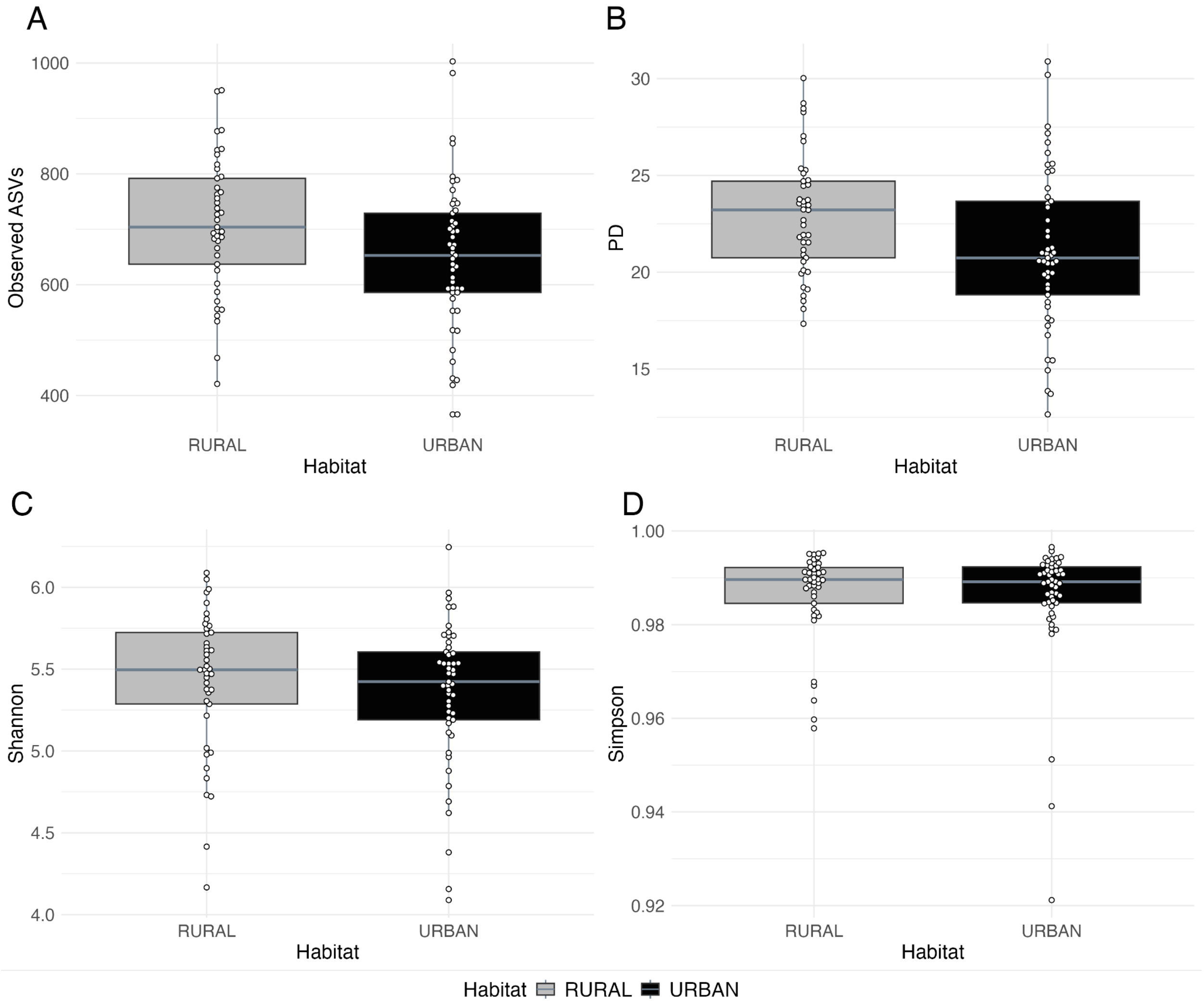


Figure S3. Boxplots of measures of alpha diversity (A. ASV richness, B. Faith's Phylogenetic Diversity, C. Shannon Diversity Index, D. Simpson Diversity Index), separated by habitat.

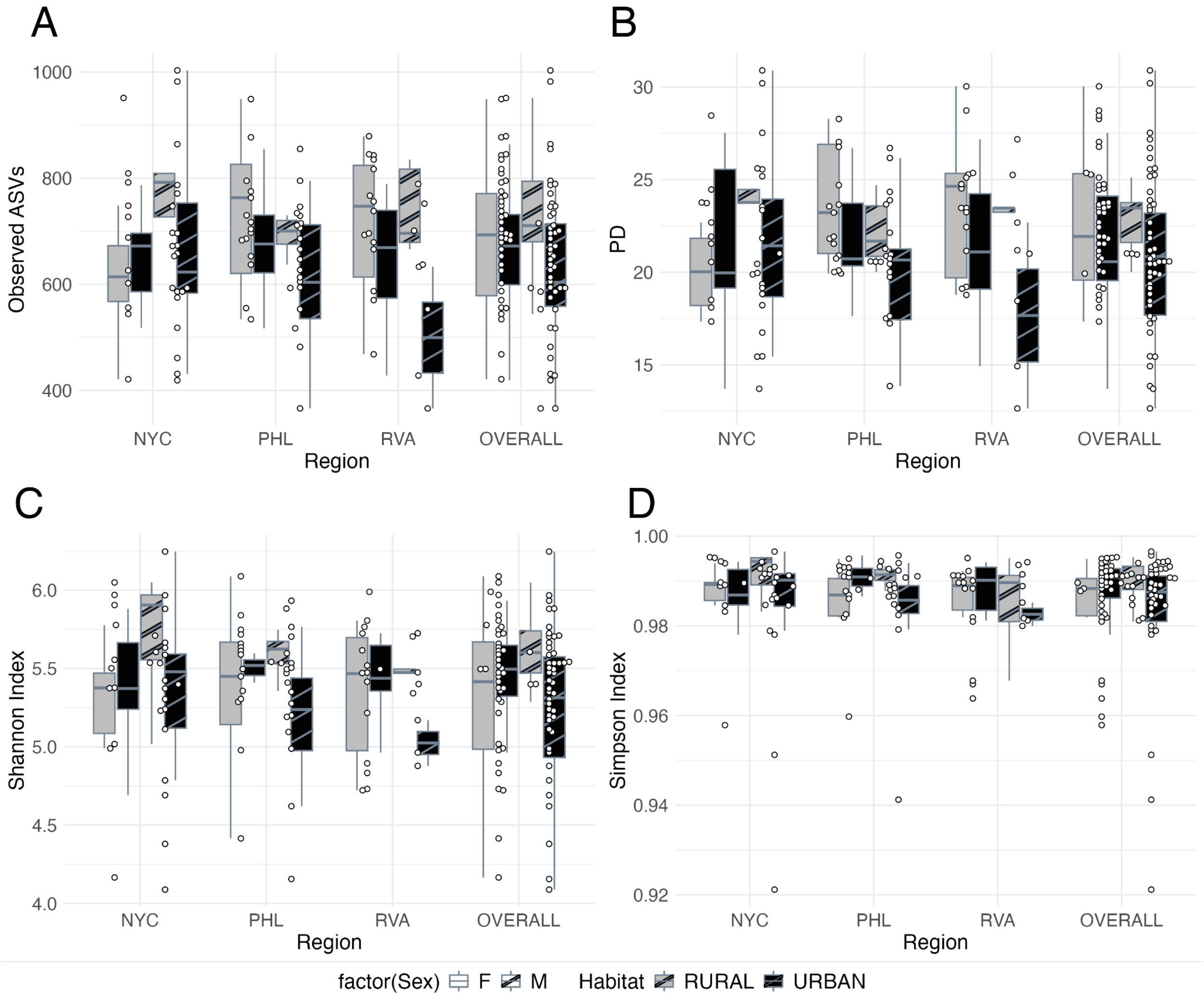


Figure S4. Boxplots of measures of alpha diversity (A. ASV richness, B. Faith's Phylogenetic Diversity, C. Shannon Diversity Index, D. Simpson Diversity Index), separated by metro region, habitat, and sex (female (solid), male (stripes)).

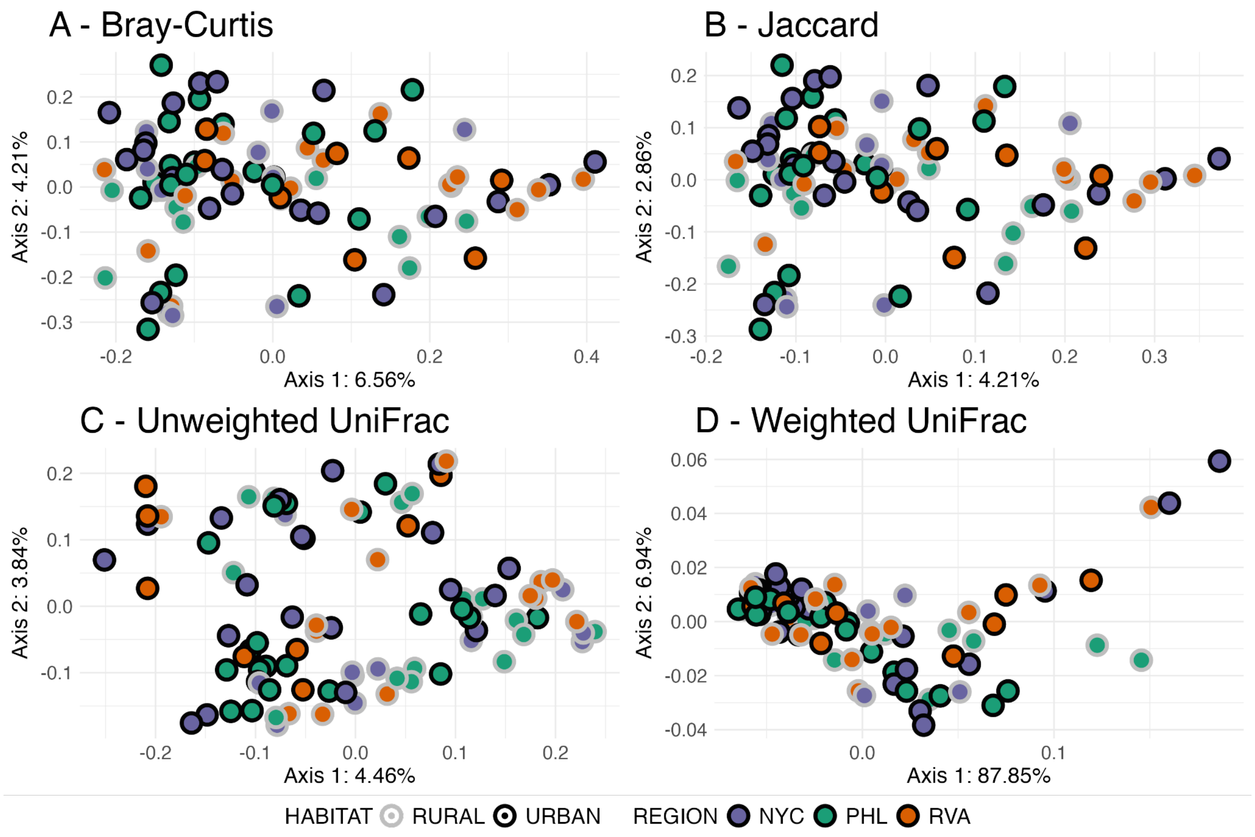


Figure S5. Samples plotted along axes 1 and 2 of Principal Coordinates Analyses (PCoA) performed using A. Bray-Curtis Distance, B. Jaccard Distance, C. Unweighted UniFrac Distance, and D. Weighted UniFrac Distance. Outer circles of points distinguish habitat and fill colors distinguish regions.

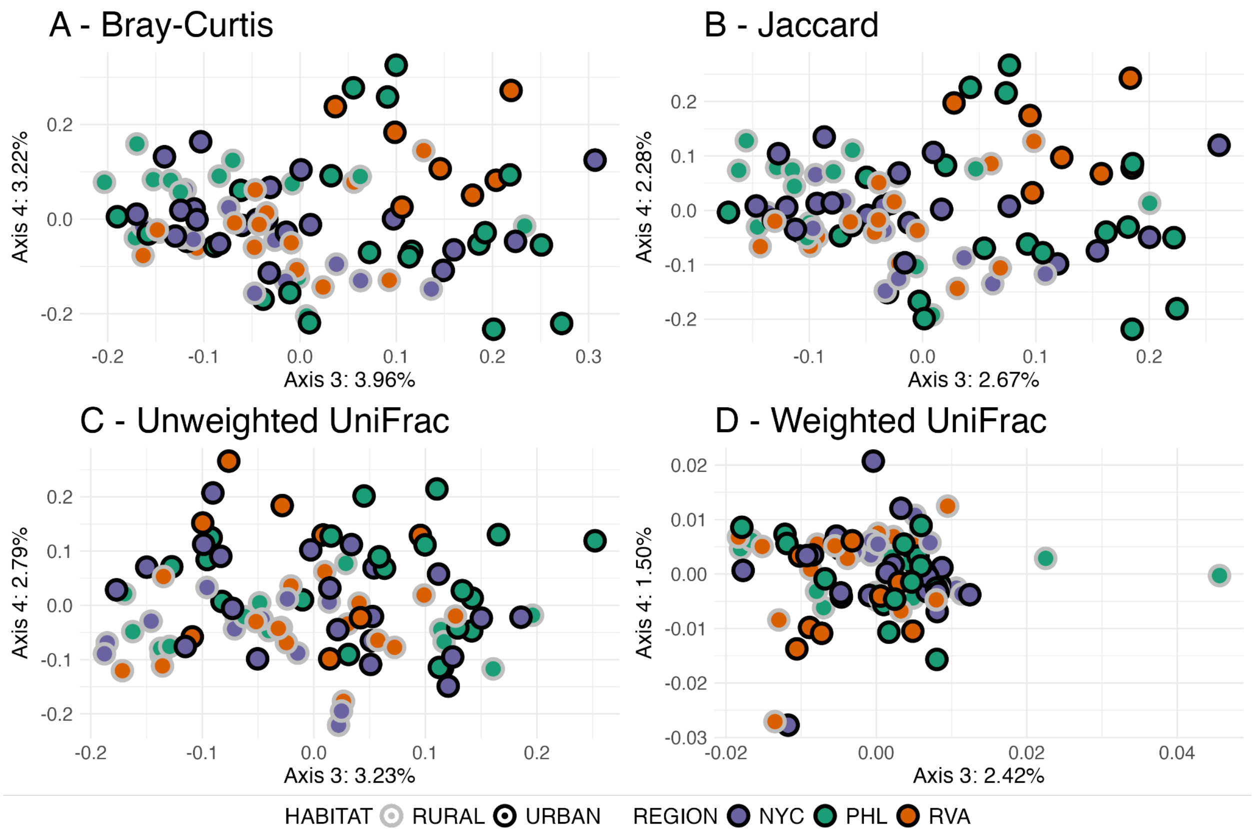


Figure S6. Samples plotted along axes 3 and 4 of Principal Coordinates Analyses (PCoA) performed using A. Bray-Curtis Distance, B. Jaccard Distance, C. Unweighted UniFrac Distance, and D. Weighted UniFrac Distance. Outer circles of points distinguish habitat and fill colors distinguish regions.

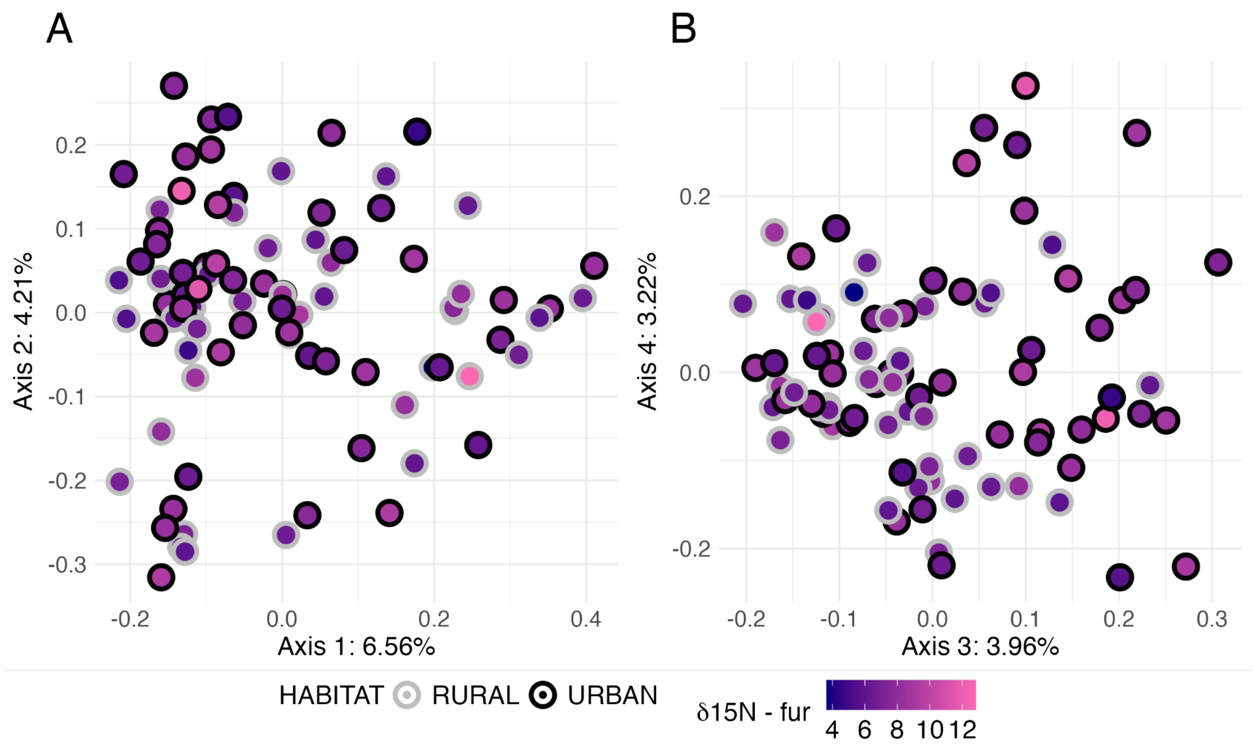


Figure S7. Samples plotted along axes a) 1 and 2, and b) 3 and 4 of Principal Coordinates Analyses (PCoA) performed using Bray-Curtis Distance. Outer circles of points distinguish habitat and fill colors distinguish stable nitrogen isotope ratios (navy: low; pink: high)

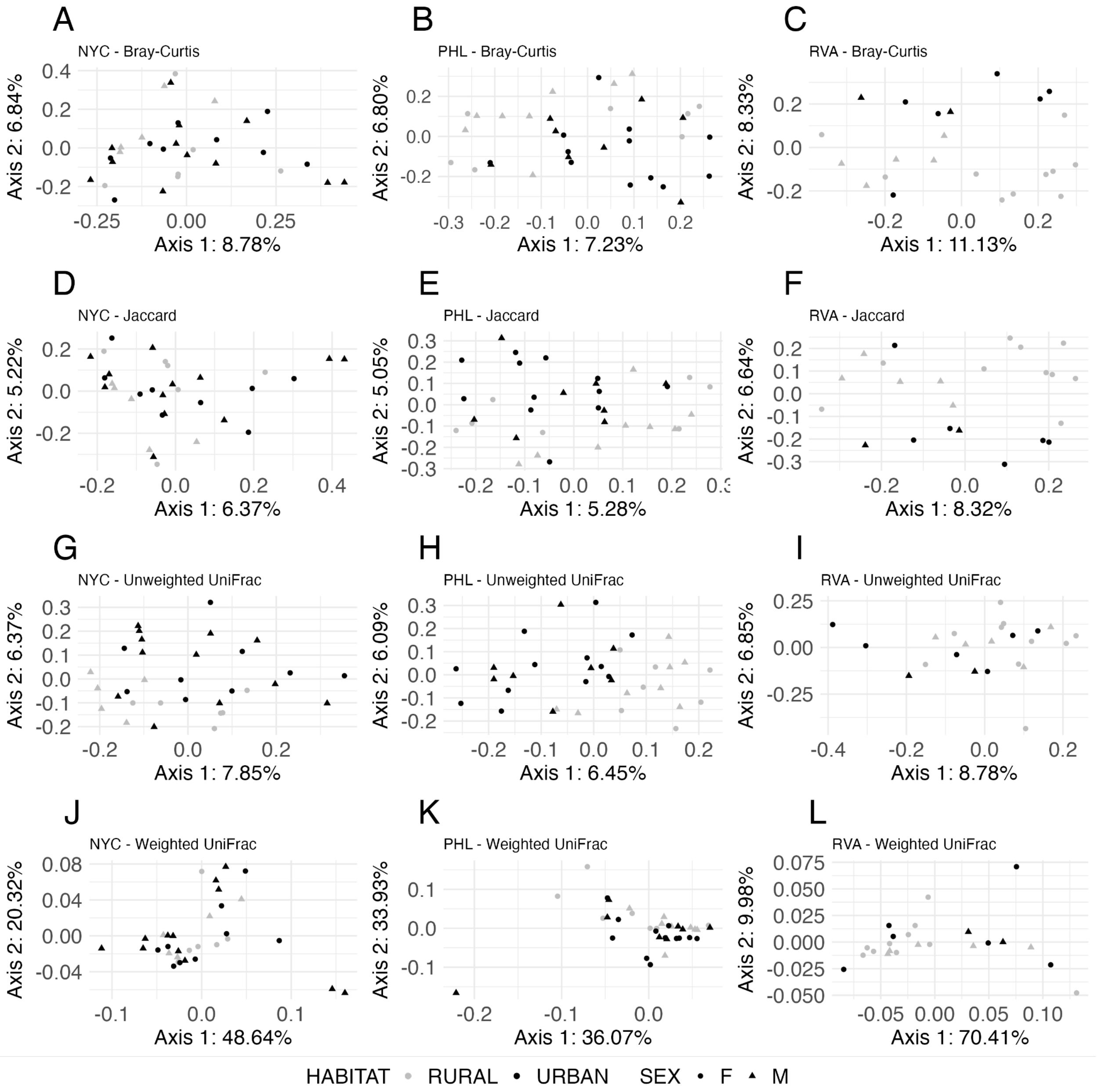


Figure S8. Samples plotted along the first two axes from Principal Coordinates Analyses (PCoAs) of A-C) Bray-Curtis Distance (BCD), D-F) Jaccard Distance, G-I) Unweighted UniFrac Distance, and J-L) Weighted UniFrac Distance between mouse gut samples from three metro regions: New York (NYC): A, D, G, & J; Philadelphia (PHL): B, E, H, & K; Richmond (RVA): C, F, I, & L. Black points refer to urban samples and gray points refer to rural samples. Circles indicate females, while triangles indicate males.

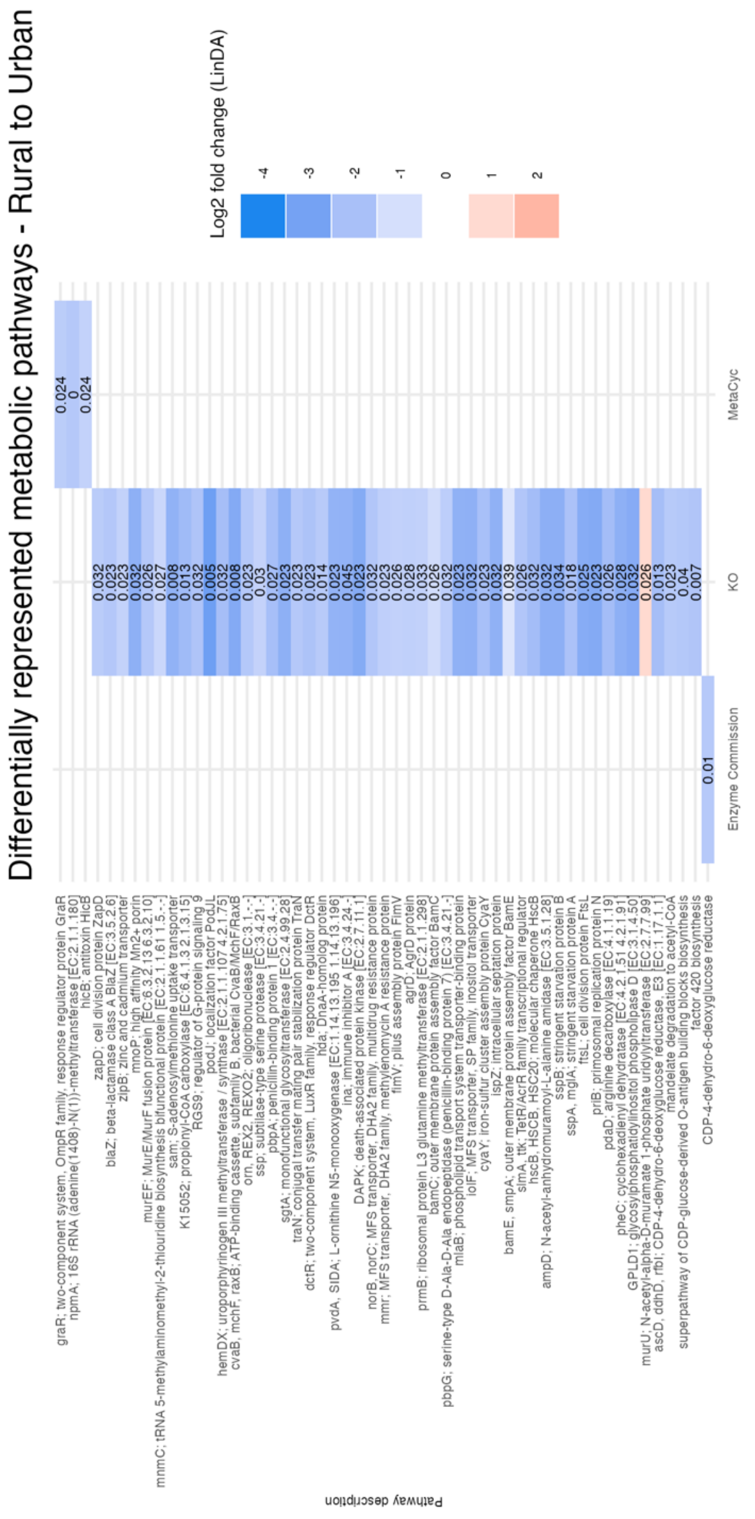


Figure S9. Heatmap of differentially represented metabolic pathways inferred from 16S sequences using the Enzyme Commission (EC), KEGG Orthologs (KO), and MetaCyc databases, respectively and the LinDA method for differential abundance. q-values are presented in tiles for each differentially represented pathway.

### Supplementary Analyses

**Pathway representation across habitat**

We also used PERMANOVA analyses to compare overall pathway representation between metadata categories, regressing distance matrices constructed from feature tables of metabolic pathways using the general formula and using the general formula: [distance matrix] ~ habitat x region x sex. Finally, we considered functional diversity and redundancy in samples, characterized here as the count of inferred KEGG orthologs and KEGG ortholog count divided by Faith's PD, respectively. These two terms were each regressed in ANOVAs against habitat, metro region, sex, and their interactions.

Despite individual metabolic pathways showing significant differential representation across habitats, there was no evidence that habitat contributed significantly to variation in overall function of gut microbiomes. PERMANOVA analyses regressing each respective picrust2-generated feature table against habitat, metro region, sex, and their interactions detected no significant contribution to variation from any of those factors (Supplementary Table 14). An ANOVA regressing diversity of inferred functions against habitat, region, sex, and their interactions identified only sex as a significant contributing factor to variation ($p=2.99 \times 10^{-2}$; Supplementary Table 15). Using Tukey's HSD, we found that the gut microbiomes of males had significantly higher functional diversity than females ($p=3.21 \times 10^{-2}$; Supplementary Table 16). Similarly, we found that sex was the only factor with evidence of contributing to variation functional redundancy (here, calculated as the ratio of a count of inferred functions and Faith's PD; $p=2.13 \times 10^{-2}$; Supplementary Table 15). Again, using Tukey's HSD, we found that males had significantly higher functional redundancy ($p=2.30 \times 10^{-2}$; Supplementary Table 16).
